# Radiosynthesis and Preclinical Evaluation of [^68^Ga]Ga-NOTA-Folate for PET Imaging of Folate Receptor β Positive Macrophages

**DOI:** 10.1101/2020.05.18.102483

**Authors:** Olli Moisio, Senthil Palani, Jenni Virta, Petri Elo, Heidi Liljenbäck, Tuula Tolvanen, Meeri Käkelä, Maxwell G. Miner, Erika Atencio Herre, Päivi Marjamäki, Tiit Örd, Merja Heinäniemi, Minna Kaikkonen-Määttä, Fenghua Zhang, Madduri Srinivasarao, Juhani Knuuti, Philip S. Low, Antti Saraste, Xiang-Guo Li, Anne Roivainen

**Author notes:** Equal contribution (Olli Moisio, Senthil Palani). Correspondence: Anne Roivainen, Turku PET Centre, Kiinamyllynkatu 4-8, FI-20520 Turku, Finland.

## Abstract

Folate receptor β (FR-β) is one of the markers expressed on macrophages and a promising target for imaging of inflammation. Here, we report the radiosynthesis and preclinical evaluation of [^68^Ga]Ga-NOTA-folate (^68^Ga-FOL). First, we determined the affinity of ^68^Ga-FOL using human FR-β expressing cells. Then, we studied atherosclerotic mice with ^68^Ga-FOL and ^18^F-FDG PET/CT. After sacrifice, the tissues excised were measured with a γ-counter for *ex vivo* biodistribution. Further, the tracer distribution and co-localization with macrophages in aorta cryosections were studied using autoradiography, hematoxylin-eosin staining and immunostaining with anti-Mac-3 antibody. Specificity of ^68^Ga-FOL was assessed in a blocking study with excess of folate glucosamine. As a last step, human radiation doses were extrapolated from rat PET data. We were able to produce ^68^Ga-FOL at high radioactivity concentration, with high molar activity and radiochemical purity. The cell binding studies showed high (5.1 ± 1.1 nM) affinity of ^68^Ga-FOL to FR-β. The myocardial uptake of ^68^Ga-FOL (SUV 0.43 ± 0.06) was 20-folds lower compared to ^18^F-FDG (SUV 10.6 ± 1.8, *P* = 0.001). The autoradiography and immunohistochemistry of aorta revealed that ^68^Ga-FOL radioactivity co-localized with Mac-3-positive macrophage-rich atherosclerotic plaques. The plaque-to-healthy vessel wall ratio of ^68^Ga-FOL (2.44 ± 0.15) was significantly higher than that of ^18^F-FDG (1.93 ± 0.22, *P* = 0.005). Blocking studies verified ^68^Ga-FOL specificity to FR. As estimated from rat data the human effective dose was 0.0105 mSv/MBq. The organ with highest absorbed dose was kidney (0.1420 mSv/MBq). In conclusion, ^68^Ga-FOL is a promising new FR-β-targeted tracer for imaging macrophage-associated inflammation.

**TABLE OF CONTENT/ABSTRACT GRAPHIC:** 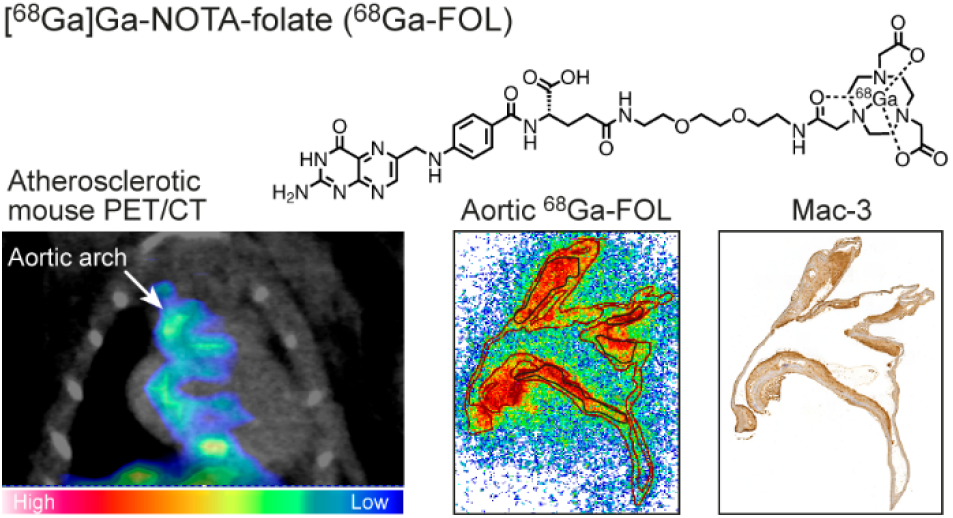

## INTRODUCTION

Folate receptor (FR) over-expression on cancer cells and during inflammation has been frequently used as a diagnostic and therapeutic tool to allow targeted delivery to tumors and inflammation^1^. The beta isoform of the folate receptor (FR-β), distinctly expressed on activated macrophages, has been recognized as a promising imaging marker for inflammatory conditions such as rheumatoid arthritis^2^. The imaging of inflammation by positron emission tomography/computed tomography (PET/CT) is currently done mainly with a glucose analog 2-deoxy-2-[^18^F]fluoro-*D*-glucose (^18^F-FDG), which reflects high consumption of glucose by macrophages and other inflammatory cell types. However, due to the non-specific nature of ^18^F-FDG, detection of inflammation adjacent to metabolically active tissues, such as the heart is difficult using ^18^F-FDG. Therefore, the development of PET tracers targeting exclusive markers are vital for the specific detection of inflammation. FR-β-targeted PET tracers investigated for this purpose so far include [^18^F]AlF-NOTA-folate (^18^F-FOL)^3–5^, [^18^F]fluoro-PEG-folate^6^ and 3’-Aza-2’-[^18^F]-fluoro-folic acid (^18^F-AzaFol)^7,8^, with the latter two already reaching initial clinical phase^9,10^. Other recently developed FR-targeted tracers include reduced ^18^F-folate conjugates^11^, ^55^Co-labeled albumin-binding folate derivatives^12^ and [^68^Ga]NOTA-folate^13^, which have been investigated for imaging FR-overexpressing tumors in preclinical studies. In a previous study, we showed that ^18^F-FOL PET successfully visualized FR-β-positive macrophages in mouse and rabbit models of atherosclerosis^3^. Atherosclerotic lesions display chronic inflammation associated with accumulation of macrophages in the affected area, providing a basis for investigating macrophage-targeted tracers.

For radiosynthesis of ^68^Ga-radiopharmaceuticals, the use of ^68^Ge/^68^Ga-generators is a common method to obtain ^68^Ga-radionuclide and can be conveniently implemented in a lab. In ^68^Ge/^68^Ga-generators, a certain amount of ^68^Ge is immobilized on a stationary phase, where the mother radionuclide decays into ^68^Ga. However, one drawback for the usage of ^68^Ge/^68^Ga-generators is that the elution capacity of ^68^Ga-radioactivity decreases as the ^68^Ge decays (physical half-life of 271 days). The reduced capacity may become an issue not only for the batch size of each production but also for the radioactivity concentration of the end product. To extent the usable lifespan of ^68^Ge/^68^Ga-generators, we have set up a remotely operated system to concentrate the ^68^Ga-eluate into small volumes from two parallel generators, which would otherwise have low individual ^68^Ga yields.

In this study, we have evaluated [^68^Ga]Ga-NOTA-folate (^68^Ga-FOL), which shares the same precursor structure as our previously studied ^18^F-FOL^3^, for imaging of inflammation. While Al^18^F labelling of NOTA-conjugates requires cyclotron facilities for the production of [^18^F]fluorine, the generator-produced ^68^Ga offers a convenient and cost-effective option for radiolabeling. First, we determined the binding affinity of ^68^Ga-FOL to human FR-β using transfected cells. Next, we investigated the uptake and specificity of intravenously (i.v) administered ^68^Ga-FOL for the detection of inflamed atherosclerotic lesions in mice and compared the tracer with ^18^F-FDG. In addition, we determined the whole-body distribution kinetics with and without blocking agent folate glucosamine in healthy rats, and estimated the human radiation dose of ^68^Ga-FOL.

## EXPERIMENTAL SECTION

### General Materials and Equipment

NOTA-folate precursor was synthesized as previously described^14^. ^68^GaCl_3_ was obtained from ^68^Ge/^68^Ga IGG-100 generators (Eckert & Ziegler, Valencia, CA, USA) via elution with 0.1 M hydrochloric acid (HCl) in water. TraceSELECT^™^ water (Honeywell, Morristown, NJ, USA) was used for radiosynthesis. Other chemicals were purchased from commercial suppliers. Chinese hamster ovary cells stably transfected with human FR-β (CHO-hFRb; CHO-FR-β^+^) were a generous gift from Philip S. Low, Purdue University, USA. FR-β negative CHO cells (CHO-FR-β^−^ control) were a generous gift from Sirpa Jalkanen, MediCity Research Laboratory, University of Turku, Finland. A dedicated small animal PET/CT (Inveon Multimodality, Siemens Medical Solutions, Knoxville, TN, USA) was used for PET/CT imaging and a gamma counter (1480 Wizard 3”, PerkinElmer/Wallac, Turku, Finland or Triathler 3”, Hidex, Turku, Finland) for radioactivity measurement of *ex vivo* tissues, blood and plasma samples. Tracer quality control and plasma metabolite analysis were performed with a LaChrom high-performance liquid chromatography (HPLC) system (Hitachi; Merck, Darmstadt, Germany) equipped with a Radiomatic 150TR flow-through radioisotope detector (Packard, Meriden, CT, USA) (radio-HPLC). Photomicroscopy images were taken with a digital slide scanner (Pannoramic 250 Flash or Pannoramic P1000, 3DHistec Ltd., Budapest, Hungary).

### ^68^Ga-FOL Radiosynthesis

#### Method 1

A fraction of ^68^Ga-eluate (0.5-1.0 mL) was mixed with an aqueous solution of 2-[4-(2-hydroxyethyl)piperazin-1-yl]ethanesulfonic acid (HEPES, 50–100 μL at a concentration of 1.2 g/mL). Then, NOTA-folate precursor (10-20 nmol in 20-40 μL water) was added, vortexed and incubated for 10 minutes at 80 °C. The mixture was then cooled down and brought to a pH of ~ 6.5 by adding 55 μL of 1 M sodium hydroxide (NaOH). The product was used without further purification. Radiochemical purity was analyzed with HPLC. The radio-HPLC conditions were as follows: 150 × 4.6 mm Jupiter 5μ C18 300 Å column (Phenomenex, Torrance, CA, USA); flow rate =□1 mL/min; wavelength λ = 220 nm; solvent A =□0.1% trifluoroacetic acid (TFA) in water; solvent B□= 0.1% TFA in acetonitrile; gradient: during 0–14 min 3% B to 25% B; 14–15 min from 25% B to 3% B.

#### Method 2

In a typical synthesis, one or two IGG-100 ^68^Ge/^68^Ga-generators were eluted with 0.1 M HCl (7.0 mL/generator) through a Strata SCX cartridge into a waste container. The cartridge was then eluted with 600 μL of 1.0 M sodium chloride/0.1 M HCl solution, from which 550 μL was transferred to the reaction vial preloaded with a mixture of HEPES (66 mg) and NOTA-folate (5–20 nmol) in 50 μL of water. Gentisic acid (10 μL, 0.1 M in water) was added to avoid radiolysis. The reaction mixture was then incubated at 40 °C for 10 minutes. The mixture was cooled down after incubation and brought to a pH of 5.5–6.5 by an addition of 30 μL of 2 M NaOH. Radiochemical purity was analyzed with HPLC similarly as described in Method 1, as well as with instant thin-layer chromatography (iTLC) (Figure S1). A 1.0 μL sample of the end product or reaction mixture was applied to a silica gel-based iTLC strip (iTLC-SG; Agilent, Santa Clara, CA, USA) and developed with 50 mM citric acid. Unbound ^68^Ga migrated up with the mobile phase with a retention factor (R_f_) of 0.8–1.0, while ^68^Ga-FOL remained at the application point (R_f_ = 0). To measure the unbound and tracer bound ^68^Ga fractions, the strip was cut into two pieces from the middle line between the baseline and the solvent front, and each piece was measured separately in a gamma counter.

Lipophilicity of ^68^Ga-FOL (distribution coefficient Log*D*) was determined as previously described^15^. To evaluate ^68^Ga-FOL stability in the injectable formulation, we kept the end product at room temperature (RT) and took samples for radio-HPLC analysis at time intervals of up to 3 hours.

### Quantification ^68^Ga-FOL Binding Affinity to FR-β

Binding specificity of ^68^Ga-FOL to FR-β was evaluated using CHO-FR-β^+^ and CHO-FR-β^−^ cells (control). The cells were cultured in growth medium - Roswell Park Memorial Institute (RPMI) 1640 medium (Gibco-Thermofisher Scientifc, Waltham, MA, USA) with 10% fetal bovine serum (FBS; Biowest, Nuaillé, France) at 37□°C in a CO_2_ incubator.

To verify FR-β expression, the cultured cells were harvested and incubated with either fluorescein isothiocyanate (FITC)-conjugated anti-human FR-β antibody (m909^16^, a gift from Philip S. Low) or allophycocyanin (APC)-conjugated anti-human FR-β antibody (mouse IgG2a, BioLegend, San Diego, CA, USA), and isotype controls (mouse IgG-FITC, mouse IgG2a-APC; BioLegend). Then, the cells were fixed using paraformaldehyde and analyzed using a fluorescence-activated cell sorting device (FACS, Fortessa flow cytometer; BD Biosciences, Franklin Lakes, NJ, USA) and Flowing software (Cell Imaging and Cytometry Core, Turku Bioscience, Turku, Finland).

After verifying the presence of FR-β on the cells, either CHO-FR-β^+^ or CHO-FR-β^−^ cells were cultured on one side of a 92-mm petri dish in a tilted position (manufacturer’s protocol, Ridgeview for ligand binding studies) in growth medium at 37□°C in a CO_2_ incubator. The other side of the petri dish with no cells was used as background control for non-specific binding of ^68^Ga-FOL. Once the cells attained a confluent monolayer, the growth medium from the petri dishes was removed and phosphate-buffered saline (PBS) containing calcium and magnesium with 10% FBS (binding medium) was added and the cells were incubated at 37□°C in a CO_2_ incubator for 30 minutes, to starve the cells of folate. After incubation, the cells were rinsed with binding medium (2 × 2 mL). A LigandTracer Yellow instrument (Ridgeview Instruments AB, Uppsala, Sweden) was then used to measure the dissociation constant (K_*D*_) for ^68^Ga-FOL. The assay protocol with LigandTracer Yellow contains consecutive radioactivity measurements of the target (cell region) and of the opposite background (no cell region on the petri dish).

Radioactivity was measured in each region for 30 seconds as raw counts per second (cps) with a delay of 5 seconds over the time course of the experiment. The target regions (cps) were corrected for background signal and for radioactive decay. To detect the background radioactivity or noise picked by the instrument, 5 mL of binding medium was added to the cells on the petri dish. After 15 minutes, ^68^Ga-FOL was added stepwise to achieve concentration range of 1 nM to 80 nM, followed by replacement with fresh binding medium to measure the dissociation. The ratio of bound ^68^Ga-FOL (to the cells) to background (petri dish) and K_*D*_ were calculated with TraceDrawer software (Ridgeview Instruments AB).

### Animal Experiments

Low-density lipoprotein receptor deficient mice expressing only apolipoprotein B100 (LDLR^−/−^ApoB^100/100^, strain #003000, The Jackson Laboratory, Bar Harbor, ME, USA) were used to induce atherosclerosis. The mice were fed with high-fat diet (HFD; 0.2% total cholesterol, TD 88137, Envigo, Madison, WI, USA) starting at the age of 2 months and maintained for 3-5 months. C57BL/6JRj mice (Central Animal Laboratory of the University of Turku) fed with a regular chow diet were used as healthy controls. In total, 17 LDLR^−/−^ ApoB^100/100^ (34.7 ± 5.5 g) and 6 healthy control mice (29.65 ± 1.9 g) were studied. In addition, 6 Sprague-Dawley rats (135.9 ± 17.1 g) from the Central Animal Laboratory of the University of Turku were studied.

All animals were housed at the Central Animal Laboratory of the University of Turku and had *ad libitum* access to water and food throughout the study. All animal experiments were approved by the national Animal Experiment Board in Finland (license number ESAVI/4567/2018) and were carried out in compliance with European Union directive (2010/63/EU).

### Mouse Studies

#### PET/CT Imaging

The mice were fasted for 4 hours prior to imaging, anesthetized with isoflurane (4–5% induction, 1–2% maintenance), and placed on a heating pad. Then, mice were i.v. administered with ^18^F-FDG (14.4 ± 0.2 MBq) via a tail vein cannula and again on the following day with ^68^Ga-FOL (20.1 ±1.0 MBq). Immediately after PET, an iodinated contrast agent (100 μL of eXIATM160XL, Binitio Biomedical Inc., Ottawa, ON, Canada) was i.v. injected and a high-resolution CT was performed for anatomical reference. To analyze PET/CT images, we used the free Carimas 2.10 software (Turku PET Centre, Turku, Finland, www.turkupetcentre.fi/carimas/). We defined regions of interest (ROIs) for the myocardium in coronal PET/CT images, using the contrast-enhanced CT as an anatomical reference as previously described^2^. The results were expressed as normalized for the injected radioactivity dose and animal body weight, i.e. as standardized uptake values (SUVs).

#### Ex Vivo Biodistribution

To study the specificity of ^68^Ga-FOL uptake, an *in vivo* blocking study was performed with another group of HFD-fed LDLR^−/−^ApoB^100/100^ mice i.v. administered with ^68^Ga-FOL alone or with 100-fold molar excess of folate glucosamine together with ^68^Ga-FOL. Mice were i.v. injected with ^68^Ga-FOL (11.3 ± 0.8 MBq) and euthanized after 60 minutes. Various tissues were excised and weighed, and their radioactivity was measured with a γ-counter (Triathler 3”, Hidex, Turku, Finland). After compensating for radioactivity remaining in the tail and cannula, the *ex vivo* biodistribution of ^68^Ga-radioactivity results were expressed SUV, and blocking *vs.* non-blocking results were compared.

#### Autoradiography, Histology and Immunostainings

Following PET/CT imaging, the dissected aortic arch was prepared into 20 μm and 8 μm cryosections. The 20 μm cryosections were used for digital autoradiography analysis as previously described^3^. Briefly, the sections were apposed on an Imaging Plate BAS-TR2025 (Fuji, Tokyo, Japan), and the plates were subsequently scanned on Fuji Analyzer BAS-5000 (Fuji, Tokyo, Japan) after an exposure time of 3 hours for ^68^Ga-FOL and at least 4 hours for ^18^F-FDG. After the scanning, sections were stored at −70°C until staining with hematoxylin and eosin (H&E) and scanned with a Pannoramic digital slide scanner. Autoradiographs were analyzed using Tina 2.1 software (Raytest Isotopemessgeräte, GmbH, Straubenhardt, Germany) and the uptake of ^68^Ga-FOL and ^18^F-FDG was corrected for injected radioactivity dose per unit body mass and radioactive decay during exposure and expressed as photostimulated luminescence per square millimeter (PSL/mm^2^). For immunohistochemistry, adjacent 8 μm sections were used to investigate co-localization of ^68^Ga-FOL with Mac-3-positive macrophages. The sections were incubated with anti-mouse Mac-3 antibody (1:1000, BD Biosciences, Franklin Lakes, NJ, USA) and a color reaction was subsequently developed using 3.3’-diaminobenzidine (Bright-DAB, BS04-110).

#### In Vivo Stability

To determine the *in vivo* stability of ^68^Ga-FOL, plasma samples collected from atherosclerotic mice (*n* = 3) at 60 minutes post-injection were analyzed using radio-HPLC. Blood samples were collected in heparinized tubes and centrifuged in 4 ^o^C for 5 minutes at 2,118 × *g*. Plasma proteins were precipitated with 10% sulfosalicylic acid (1:1 *v/v*) followed by centrifugation for two minutes at 14,000 × *g* at RT. The supernatant was analyzed with radio-HPLC. Standard samples were prepared by adding ^68^Ga-FOL tracer to 500 μL of plasma supernatant collected from mice without the administration of tracer. Both standard and metabolite samples applied to radio-HPLC analysis were normalized to a final volume of 1 mL by dilution with radio-HPLC solvent A if necessary. The radio-HPLC conditions were as follows: 250 × 10 mm Jupiter Proteo 5μ C18 90 Å column (Phenomenex, Torrance, CA, USA); flow rate = □5 mL/min; solvent A = □ 0.1% TFA in water; solvent B = 0.1% TFA in acetonitrile; gradient was during 0-11 min from 3% B to 25% B, during 11–12 min from 25% B to 100% B, during 12–14 min 100% B.

### Rat Studies

In order to determine distribution kinetics and estimate human radiation dose, a dynamic whole-body ^68^Ga-FOL PET/CT was performed in six healthy rats. In addition, three of the rats were also subjected to a blocking experiment with a co-injection of a 100-fold excess of folate glucosamine. The rats were injected with 10.3 ± 0.4 MBq of ^68^Ga-FOL and PET imaged for 60 minutes. After the imaging, the rats were euthanized, various tissues excised, weighed and measured for radioactivity and plasma samples were analyzed using radio-HPLC as described above. Using CT as an anatomical reference, quantitative PET image analysis was performed by defining ROIs on the main organs and time-activity curves were extracted with Carimas software. Human radiation dosimetry was estimated from the rat data using the OLINDA/EXM 2.2 software^17^.

### Statistical Analysis

Results are presented as mean ± SD. Differences between groups were analyzed by the unpaired Student *t*-test using Microsoft Excel. *P* values of less than 0.05 were considered statistically significant.

## RESULTS

### Radiosynthesis

#### Method 1

^68^Ga-FOL was produced with 345.8 ±118.9 MBq/mL (*n* = 13) radioactivity concentration and high radiochemical purity (96.6 ± 2.7%). The molar activity was 21.8 ± 6.9 GBq/μmol at the end of radiosynthesis. Distribution coefficient Log*D* of ^68^Ga-FOL was −3.28 ± 0.33 (*n* = 3), indicating high hydrophilicity.

#### Method 2

We set up a straightforward system using commonplace hospital pressure IV-tubing (Argon, Frisco, TX, USA) and one-way check valves (B. Braun, Melsungen, Germany) and a 3-way valve (Bürkert, Huntersville, USA) to allow the elution of two generators through an SCX cartridge in a remotely operated manner. The system is outlined in Figure 1A. When two parallel ^68^Ge/^68^Ga-generators (aged 16 and 20 months with nominal radioactivities of 1.85 GBq and 2.20 GBq at the time of manufacture, respectively) we were able to produce 492.9 ± 24.2 MBq (*n* = 5) of ^68^Ga-FOL at high radioactivity concentration (782.5 ± 38.4 MBq/mL) and radiochemical purity (99.3 ± 0.2%). The molar activity was 49.3 ± 2.42 GBq/μmol.

**Figure 1.**
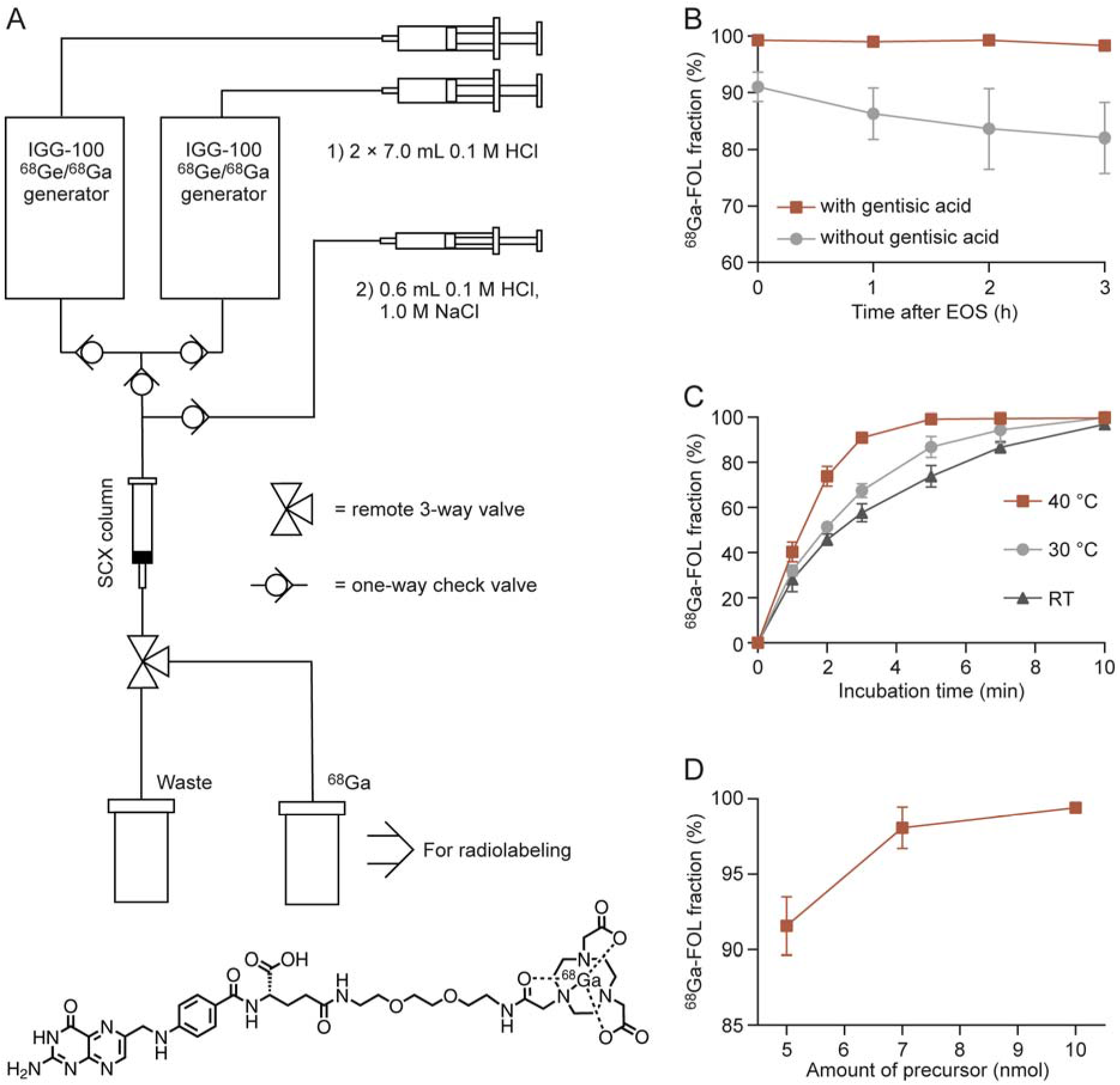
(A) The remotely operated system for the concentration of ^68^GaCl_3_ eluted from two parallel generators and a flow chart for ^68^GaCl_3_ elution and concentration. The molecular weight of ^68^Ga-FOL is 908.82 Da. (B) The effect of gentisic acid on the stability of the end product as determined with radio-HPLC (*n* = 3, reaction conditions: 40 °C, 15 nmol of precursor). (C) ^68^Ga-FOL fraction of total radioactivity during incubation, measured with iTLC-SG (*n* = 3, reaction conditions: 15 nmol of precursor, 10 μL of 0.1 M gentisic acid). (D) Effect of precursor amount on radiochemical purity at the end of synthesis (EOS) as measured with iTLC-SG (*n* = 3, reaction conditions: 40 °C, 10 μL 0.1 M gentisic acid).

### *In Vitro* Quantification of ^68^Ga-FOL Binding Affinity to FR-β

FACS analyses verified that FR-β was clearly expressed on CHO-FR-β^+^ cells but not on CHO-FR-β^−^ cells (Figure 2A-C). In binding assays, we found that with a stepwise increase in concentration of ^68^Ga-FOL from 1 nM to 80 nM, the binding of ^68^Ga-FOL to CHO-FR-β^+^ cells was gradually increased, and showed a K*D* of 5.1 ± 1.1 nM (*n* = 3). Whereas there was no clear accumulation of ^68^Ga-FOL to CHO-FR-β^−^ cells even at concentrations up to 40 nM (Figure 2D).

**Figure 2.**
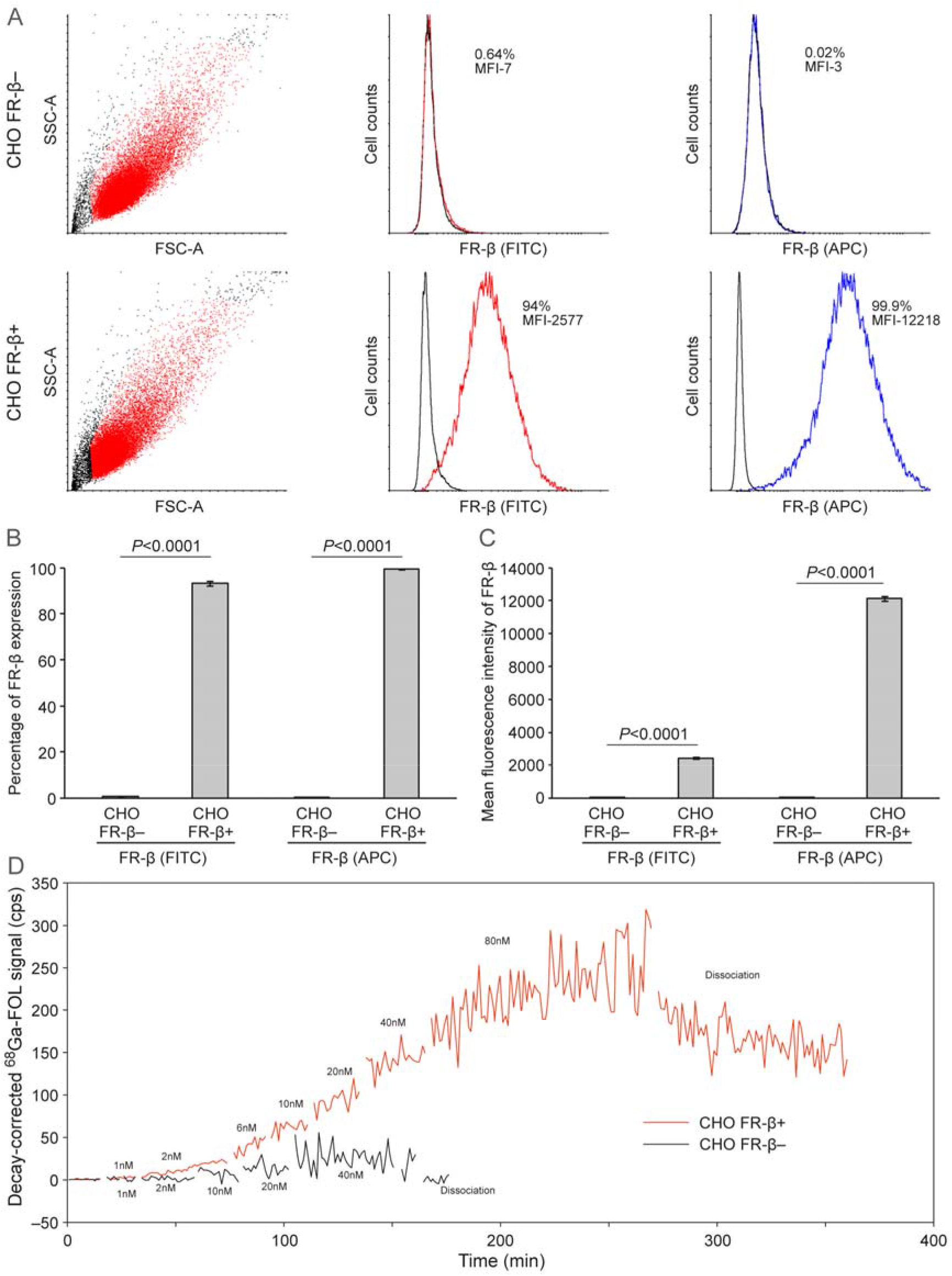
(A) Representative figures of flow cytometric analyses showing FR-β expression on human FR-β-negative and FR-β-positive CHO cells stained with FITC-conjugated anti-FR-β antibody (red) or APC-conjugated anti-FR-β antibody (blue) and isotope controls (black). Quantification of FR-β expression presented as (B) percentage and (C) mean fluorescence intensity. (D) Representative real time binding affinity of ^68^Ga-FOL measurement with LigandTracer. The graph generated with TracerDrawer, shows raw counts per second (cps) after corrected for background signal and for radioactive decay.

### ^68^Ga-FOL Detects Macrophage-rich Lesions in Atherosclerotic Mice

We evaluated the biodistribution of i.v. administered ^68^Ga-FOL in mice using *in vivo* PET/CT, *ex vivo* gamma counting of excised tissues and *ex vivo* autoradiography of aorta cryosections. To study the specificity of ^68^Ga-FOL to FR-β, blocking with co-injection of a molar excess of folate glucosamine was performed. In addition, ^68^Ga-FOL was compared with ^18^F-FDG in a head-to-head PET/CT imaging setting and by *ex vivo* autoradiography.

The *in vivo* stability of ^68^Ga-FOL was good; 60 minutes after i.v. injection, the amount of intact tracer was 63.3 ± 1.2% of the total plasma radioactivity in LDLR^−/−^ApoB^100/100^ mice (*n* = 3, Figure S2).

Our *ex vivo* results revealed that the aortic uptake of ^68^Ga-FOL was higher in atherosclerotic mice (SUV 0.75 ± 0.12) than in healthy controls (SUV 0.41 ± 0.10, *P* = 0.004) or atherosclerotic mice from the blocking study (SUV 0.09 ± 0.03, *P* = 0.001). Further, the atherosclerotic aorta radioactivity concentration was 3-fold greater than that of blood (SUV 0.23 ± 0.09). The highest radioactivity uptake was seen in FR-positive kidneys^18^ in both atherosclerotic and control mice (SUV 22.30 ± 3.28 and 20.27 ± 5.48, respectively, *P* = 0.49) and it was significantly reduced in the blocking study of atherosclerotic mice (SUV 2.65 ± 1.80, *P* = 0.0002). The radioactivity of other tissues was much lower than that of the kidneys. Besides kidneys, the folate glucosamine blocking in atherosclerotic mice reduced the radioactivity concentration in many other tissues too, importantly in the aorta by 88% (Supporting Information Table S1).

A comparison of the two tracers by *in vivo* PET/CT revealed that ^68^Ga-FOL uptake in myocardium (SUV 0.43 ± 0.06) was significantly lower than that of ^18^F-FDG (SUV 10.6 ± 1.88, *P* = 0.001, Figure 3A,B).

**Figure 3.**
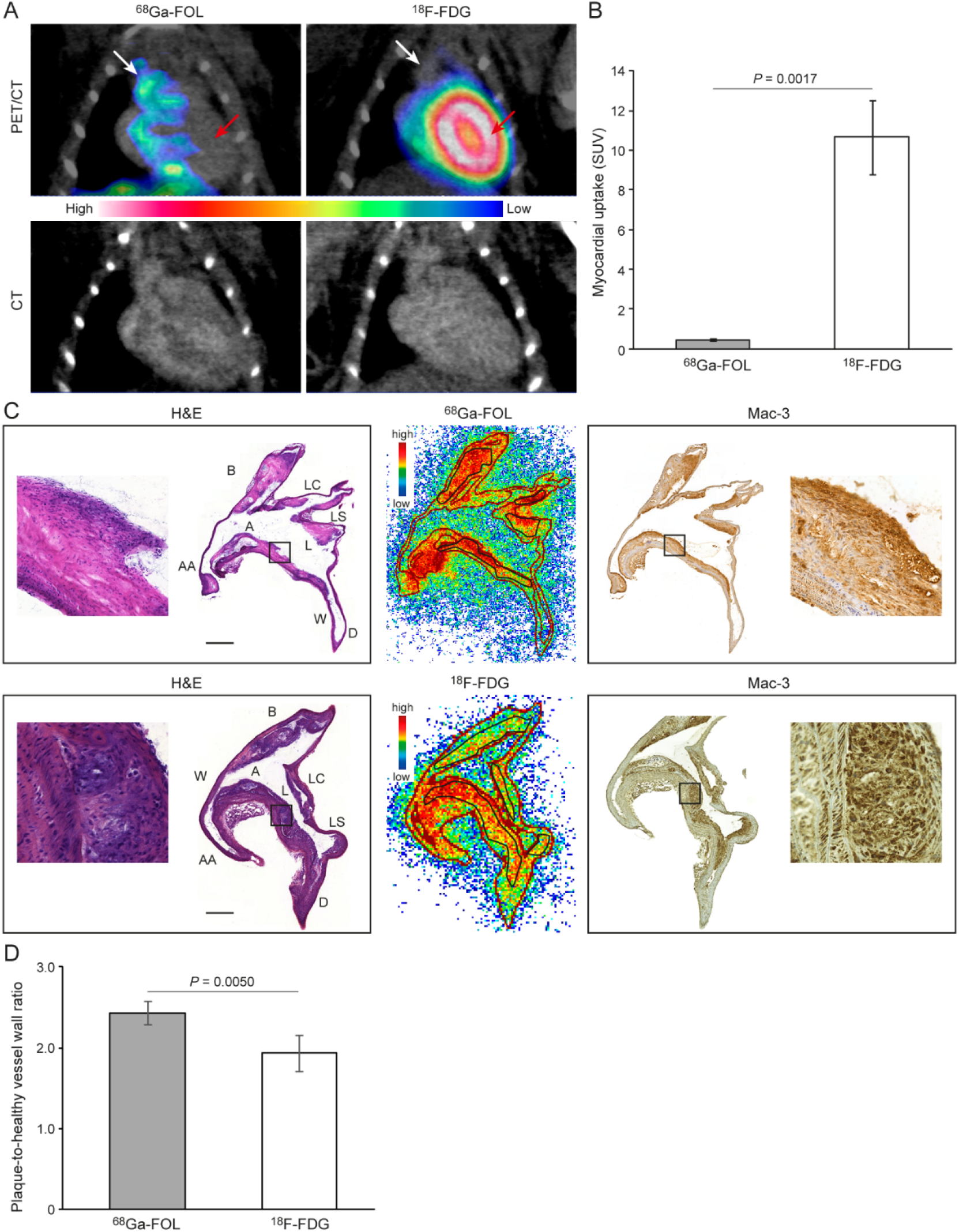
(A) Coronal PET/CT and CT images of atherosclerotic mouse administered with ^68^Ga-FOL or ^18^F-FDG. White arrows show aortic arch and red arrows show myocardium region, respectively. (B) Quantification of myocardial PET data showing significant difference between the tracers. (C) Hematoxylin-eosin (H&E) staining and autoradiography images from representative aorta cryosections, and Mac-3 macrophage marker staining on consecutive aorta cryoscetions, Black rectangle indicates the plaque region, which is zoomed. (D) Quantification of autoradiography ^68^Ga-FOL data showing significant difference between the groups. (E) Quantification of autoradiography data showing significant difference between the tracers. Scale bar = 0.5 mm. A = arch; AA = ascending aorta; B = brachiocephalic artery; D = descending thoracic aorta; L = lesion; LC = left common carotid artery; LS = left subclavian artery; W = wall.

In order to further elucidate ^68^Ga-FOL and ^18^F-FDG uptake in the aortas of atherosclerotic mice in more detail, we analyzed radioactivity using autoradiography and H&E staining of aortic cryosections followed by macrophage-detecting immunohistochemistry on adjacent tissue cryosections. The results revealed that ^68^Ga-FOL and ^18^F-FDG radioactivity co-localized with Mac-3-positive macrophage-rich plaques (Figure 3C). The plaque-to-healthy vessel wall ratio of ^68^Ga-FOL (2.44 ± 0.15) was significantly higher than that of ^18^F-FDG (1.93 ± 0.22, *P* = 0.005, Figure 3D).

### Distribution Kinetics in Rats and Estimate Human Radiation Dose of ^68^Ga-FOL

As in mice, ^68^Ga-FOL showed fast renal excretion and the highest uptakes in kidneys, urine, salivary glands, liver and spleen (Figure 4). Co-injection of ^68^Ga-FOL and molar excess of folate glucosamine, clearly decreased tracer uptake in several organs but increased urinary excretion.

**Figure 4.**
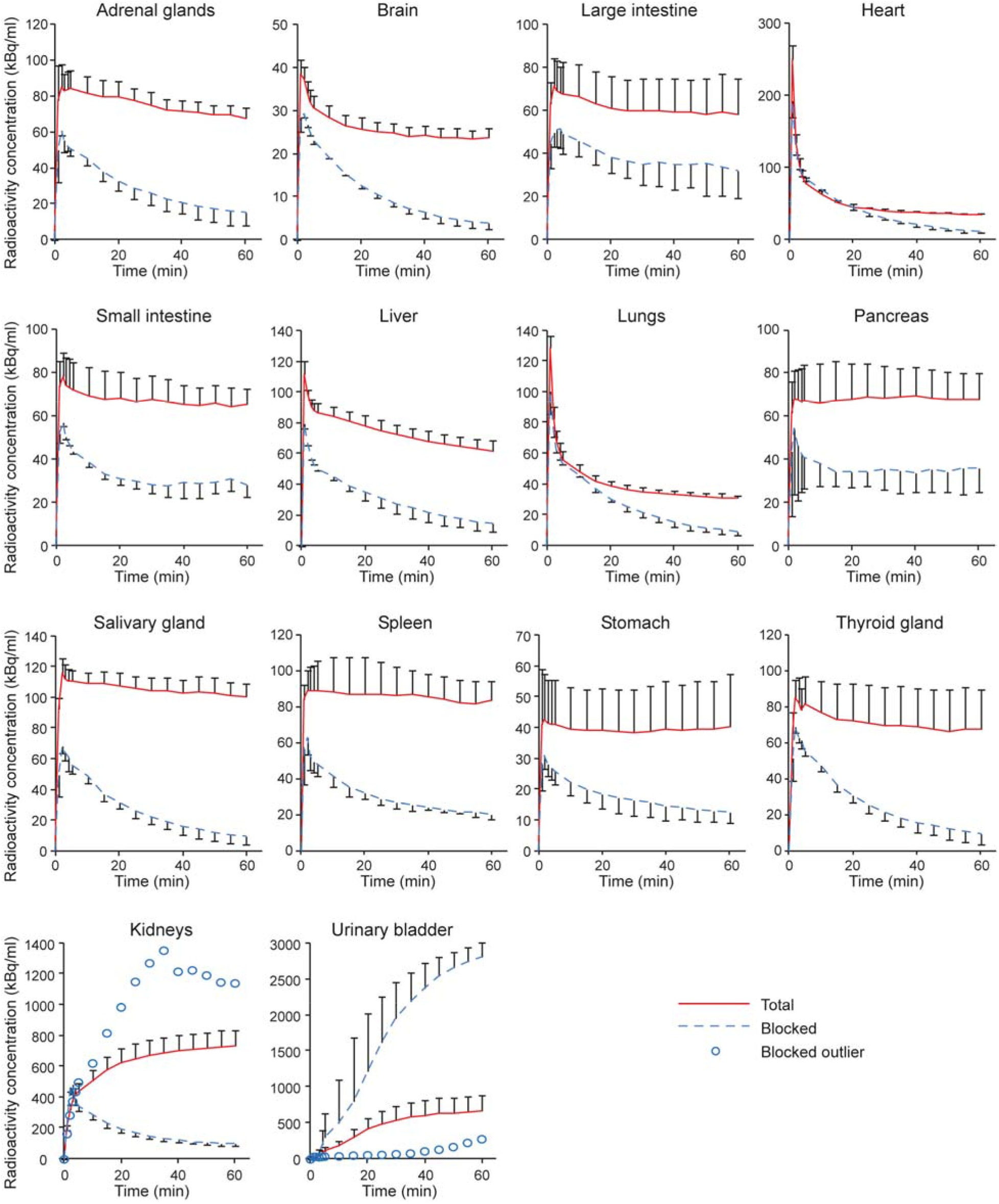
Time-activity curves of healthy rat tissues after i.v. injection of ^68^Ga-FOL or co-injection of ^68^Ga-FOL and molar excess of folate glucosamine (blocking experiment).

Radio-HPLC analysis of plasma samples revealed good *in vivo* stability of ^68^Ga-FOL (Figure S2); at 60 minutes post-injection 71.8 ± 1.5% of the total radioactivity was still counting from the intact tracer in healthy rats (*n* = 3) without blocking and 88.0 ± 0.7% (*n* = 3, *P* = 0.0002) when blocked with folate glucosamine. Extrapolating from the rat PET data, the estimated human effective dose for a 73 kg man was 0.0105 mSv/MBq. The most critical organ was the kidney (0.1420 mSv/MBq) (Supporting Information Table S2).

## DISCUSSION

In this study, we developed a new method for effective synthesis of ^68^Ga-FOL, which could be applicable to generate other ^68^Ga-tracers. We report that ^68^Ga-FOL binds to FR-β with high affinity, and when i.v. administered accumulates in atherosclerotic lesions in mice. Importantly, ^68^Ga-FOL showed lower myocardial uptake and higher plaque-to-healthy vessel wall ratio compared to ^18^F-FDG. Human radiation dose of ^68^Ga-FOL was low as estimated from the rat data.

To produce ^68^Ga-FOL, we have used a fractionation method to obtain ^68^GaCl_3_ from a ^68^Ge/^68^Ga-generator for a chelation reaction with the precursor compound NOTA-folate (Radiosynthesis, Method 1). This is a well-established method that we have used previously and total radiosynthesis takes less than 20 minutes. The radioactivity concentrations are sufficiently high for applications such as cell binding experiments. However, for *in vivo* studies in mice, it is increasingly difficult to produce sufficiently high radioactivity concentration by using fractionation-based synthesis as the generator’s ^68^Ga yield decreases over time. This has motivated us to explore alternative methods to utilize the generators in a more effective way. In our lab, we indeed have fully automated ^68^Ga-radiosynthesis devices for ^68^Ga-eluate concentration in the production of ^68^Ga-radiopharmaceuticals compliant with Good Manufacturing Practice (GMP), but it is more cost-effective to build up a simpler system for the synthesis of preclinical non-GMP tracers. Similar approaches have been reported from other labs as well^19^. Accordingly, we have used a 3-way tubing system to connect two parallel generators to a SCX-cartridge abstracting ^68^Ga-nuclide from generator eluates, and a remotely controlled 3-way valve to direct the liquids either to a waste vial or a vial for receiving concentrated ^68^Ga-radioactivity (Figure 1A). It is optional to elute ^68^Ga from both generators or from either of them, which increases flexibility. With this system we have managed to concentrate > 90% radioactivity from the generators into a volume of 0.6 mL at radioactivity concentrations up to 1 GBq/mL approximately, when using two generators at ages 16 and 20 months, respectively. This is a large improvement for radiolabeling applications but initially lead to other issues. When using concentrated ^68^Ga-eluate, we have concurrently observed a significant amount of radioactive side product as detected by radio-HPLC quality analysis of ^68^Ga-FOL. Based on our experiences in the radiosynthesis of the corresponding ^18^F-FOL^3^, we have presumed that the side product is caused by radiolysis at high radioactivity concentrations. By adding gentisic acid as a radical scavenger, the formation of the side product has been effectively prevented, the ^68^Ga-FOL being stable for at least 3 hours (Figure 1B, stability for longer time has not been tested). Furthermore, we have observed that the chelation reaction with this type of post-processed ^68^Ganuclide is rather efficient. Even at RT, the reaction can reach near completion in 10 minutes (Figure 1C) and the total radiosynthesis time is only 5 minutes longer than the synthesis with the fractionation method. In typical cases, 10 nmol of precursor is sufficient for each synthesis (Figure 1D), affording the formation of ^68^Ga-FOL with high molar activities (49.3 ± 2.4 GBq/μmol). Thus, this indicates the approach is effective for the synthesis of ^68^Ga-FOL.

Previously, ^99m^Tc-EC20 and ^111^In-EC0800, two folate-based imaging agents for single-photon emission computed tomography (SPECT) have been shown to detect atherosclerotic lesions in mice^20,21^. Additionally, PET tracer 3′-aza-2′-^18^F-fluorofolic acid has been shown to detect FR-β-positive macrophages in human atherosclerotic plaques *in vitro*^8^. In our previous studies, we have reported ^18^F-FOL specificity to FR-β-positive macrophages and detection of inflamed atherosclerotic plaques in mice and rabbits as well as in human tissue sections^3^. However, ^68^Ga-FOL binding affinity to human FR-β and ability to detect atherosclerotic lesions in comparison to ^18^F-FDG has not been evaluated before. Our *in vitro* binding assay of ^68^Ga-FOL with CHO-FR-β^+^ and CHO-FR-β^−^ cells showed good specificity and high affinity to FR-β (5.1 ± 1.1 nM), which falls close to the binding affinity of ^18^F-FOL (1.0 nM) to FR–positive tumor xenografts reported earlier^14^. Our blocking studies in mice and rats further supported tracer’s specificity to FR. Mouse studies confirmed the ability to detect macrophage-rich inflammatory lesions. The observed low myocardial uptake is beneficial for detection of atherosclerotic lesions in coronary arteries in prospective PET/CT studies of patients with coronary heart disease. When compared to our earlier ^18^F-FOL study, ^68^Ga-FOL showed similar plaque-to-healthy vessel wall ratios (2.44 ± 0.15 for ^68^Ga-FOL and 2.60□±□0.58 for ^18^F-FOL). However, the *in vivo* stability of ^68^Ga-FOL in mice (63 ± 1 % intact tracer at 60 minutes post-injection) was slightly lower compared to our previous studies with ^18^F-FOL (85 ± 6% at 60 min post-injection)^3^. The human effective dose of ^68^Ga-FOL extrapolated from rat data (0.0105 mSv/MBq) is low and within the same range as other ^68^Ga-tracers^22-24^.

In conclusion, we demonstrated the feasibility to produce ^68^Ga-FOL at high radioactivity concentration, with high molar activity and radiochemical purity by using a straightforward system for extracting ^68^Ga-nuclide from aged ^68^Ge/^68^Ga-generators. Similar method may be applicable for producing other ^68^Ga-tracers. The preclinical results of ^68^Ga-FOL are in the line with our previous studies using ^18^F-FOL and corroborate FR-β as an imaging target for detection of inflamed atherosclerotic lesions.

## ACKNOWLEDGMENTS

The authors thank Aake Honkaniemi and Timo Kattelus for technical assistance. This work was financially supported by the Academy of Finland (grant numbers 314553, 314554, 314556), Sigrid Jusélius Foundation and Jane and Aatos Erkko Foundation.

## SUPPORTING INFORMATION

Figure S1: Representative iTLC-SG autoradiographs and chromatographs; Figure S2: Representative radio-HPLC chromatograms; Table S1: *Ex vivo* biodistribution in mice; Table S2: Human radiation doses extrapolated from the rat data.

