## Supporting Information for "Radiosynthesis and Preclinical Evaluation of [^68^Ga]Ga-NOTA-Folate for PET Imaging of Folate Receptor β Positive Macrophages"

#Equal contribution

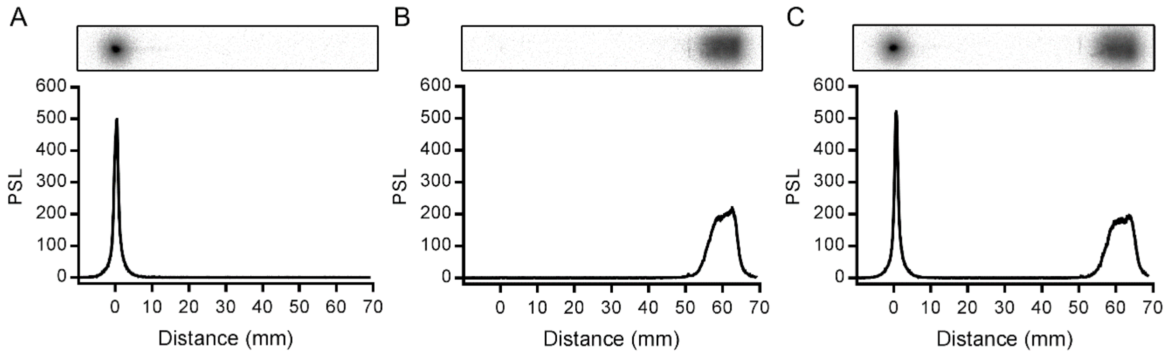

**Figure S1.** Representative autoradiographs (upper panels) and chromatograms (lower panels) of instant thin-layer thin-layer-silica gel (iTLC-SG) strips developed with 50 mM citric acid for (A) $^{68}$Ga-FOL, (B) unbound $^{68}$Ga in the reaction mixture without NOTA-folate precursor and (C) co-application of $^{68}$Ga-FOL and $^{68}$Ga.

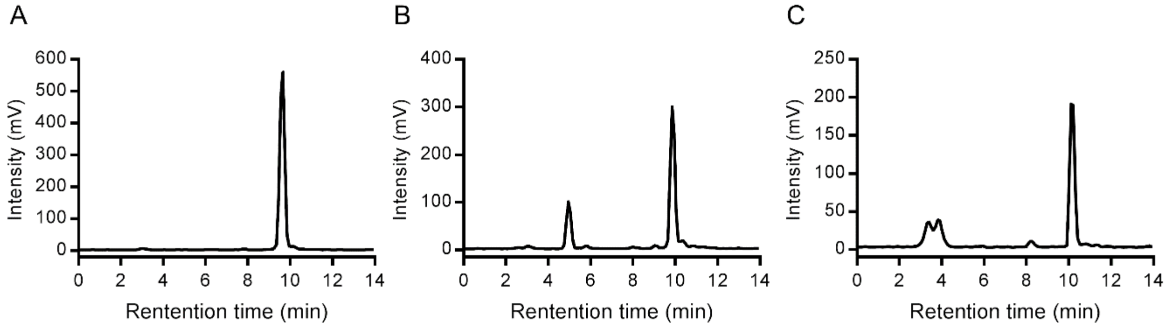

**Figure S2.** Representative radio-HPLC chromatograms of (A) $^{68}$Ga-FOL standard, (B) rat plasma at 60 minutes after $^{68}$Ga-FOL injection and (C) and mouse plasma 60 minutes after $^{68}$Ga-FOL injection.

**Table S1.** *Ex vivo* biodistribution of $^{68}$Ga-FOL at 60 minutes post-injection in mice expressed as standardized uptake values (mean ± SD)

| Tissue | LDLR$^{-/-}$ApoB$^{100/100}$ atherosclerotic mice (*n* = 4) | LDLR$^{-/-}$ApoB$^{100/100}$ atherosclerotic mice, blocking (*n* = 4) | C67BL/6JRj control mice (*n* = 6) |
|---|---|---|---|
| Aorta | 0.75 ± 0.12 | 0.09 ± 0.03 (*P* = 0.001)[a] | 0.41 ± 0.10 (*P* = 0.004)[b] |
| Brown adipose tissue | 0.29 ± 0.04 | 0.06 ± 0.02 (*P* = 0.006)[a] | 0.31 ± 0.05 (*P* = 0.51)[b] |
| Blood | 0.23 ± 0.09 | 0.29 ± 0.10 (*P* = 0.48)[a] | 0.25 ± 0.14 (*P* = 0.84)[b] |
| Bone (skull) | 0.13 ± 0.05 | 0.06 ± 0.02 (*P* = 0.04)[a] | 0.16 ± 0.04 (*P* = 0.42)[b] |
| Bone + marrow (femur) | 0.24 ± 0.18 | 0.07 ± 0.01 (*P* = 0.17)[a] | 0.18 ± 0.04 (*P* = 0.57)[b] |
| Brain | 0.08 ± 0.01 | 0.02 ± 0.00 (*P* = 0.001)[a] | 0.06 ± 0.01 (*P* = 0.02)[b] |
| Heart | 0.20 ± 0.02 | 0.05 ± 0.02 (*P*<0.0001)[a] | 0.24 ± 0.03 (*P* = 0.04)[b] |
| Intestine, small (empty) | 0.44 ± 0.06 | 0.42 ± 0.10 (*P* = 0.84)[a] | 0.47 ± 0.27 (*P* = 0.76)[b] |
| Intestine, large (empty) | 0.73 ± 0.14 | 0.19 ± 0.06 (*P* = 0.003)[a] | 0.66 ± 0.08 (*P* = 0.38)[b] |
| Kidneys | 22.30 ± 3.28 | 2.65 ± 1.80 (*P* = 0.0002)[a] | 20.27 ± 5.48 (*P* = 0.49)[b] |
| Lungs | 0.46 ± 0.03 | 0.19 ± 0.04 (*P*<0.0001)[a] | 0.39 ± 0.06 (*P* = 0.05)[b] |
| Liver | 1.04 ± 0.43 | 0.33 ± 0.13 (*P* = 0.04)[a] | 0.76 ± 0.16 (*P* = 0.29)[b] |
| Lymph node | 1.37 ± 0.57 | 0.14 ± 0.02 (*P* = 0.02)[a] | 4.07 ± 0.73 (*P* = 0.0002)[b] |
| Muscle | 0.20 ± 0.02 | 0.05 ± 0.01 (*P*<0.0001)[a] | 0.13 ± 0.02 (*P* = 0.0003)[b] |
| Pancreas | 0.43 ± 0.06 | 0.07 ± 0.02 (*P* = 0.0007)[a] | 0.37 ± 0.03 (*P* = 0.17)[b] |
| Plasma | 0.40 ± 0.15 | 0.48 ± 0.13 (*P* = 0.44)[a] | 0.34 ± 0.15 (*P* = 0.61)[b] |
| Spleen | 0.23 ± 0.07 | 0.15 ± 0.05 (*P* = 0.11)[a] | 0.28 ± 0.13 (*P* = 0.42)[b] |
| Stomach (empty) | 0.78 ± 0.14 | 0.25 ± 0.05 (*P* = 0.002)[a] | 0.65 ± 0.12 (*P* = 0.17)[b] |
| Thymus | 0.42 ± 0.13 | 0.06 ± 0.01 (*P* = 0.01)[a] | 0.33 ± 0.08 (*P* = 0.25)[b] |
| White adipose tissue | 0.22 ± 0.16 | 0.03 ± 0.01 (*P* = 0.10)[a] | 0.30 ± 0.08 (*P* = 0.40)[b] |

The blocking study was performed by injecting a 100-fold molar excess of folate glucosamine 1 minute before the $^{68}$Ga-FOL.

[a]Difference between LDLR$^{-/-}$ApoB$^{100/100}$ atherosclerotic mice *vs*. atherosclerotic mice + blocking, as assessed by independent samples *t*-test.

[b]Difference between LDLR$^{-/-}$ApoB$^{100/100}$ atherosclerotic *vs*. C57BL//6JRj control mice as assessed by independent samples *t*-test.

**Table S2.** Human dose equivalent estimates (mSv/MBq) extrapolated from the rat PET data

| Organ | Organ doses |
|---|---|
| Adrenals | 0.0243 |
| Brain | 0.0043 |
| Esophagus | 0.0121 |
| Eyes | 0.0110 |
| Gallbladder wall | 0.0139 |
| Left colon | 0.0140 |
| Small intestine | 0.0180 |
| Stomach wall | 0.0131 |
| Right colon | 0.0184 |
| Rectum | 0.0134 |
| Heart wall | 0.0078 |
| Kidneys | 0.1420 |
| Liver | 0.0118 |
| Lungs | 0.0069 |
| Pancreas | 0.0135 |
| Prostate | 0.0134 |
| Salivary glands | 0.0119 |
| Red marrow | 0.0102 |
| Osteogenic cells | 0.0096 |
| Spleen | 0.0173 |
| Testes | 0.0119 |
| Thymus | 0.0120 |
| Thyroid | 0.0113 |
| Urinary bladder wall | 0.0132 |
| Total body | 0.0130 |
| Effective dose | 0.0105 |
